# Simulation of foraging behavior using a decision-making agent with Bayesian and inverse Bayesian inference: Temporal correlations and power laws in displacement patterns

**DOI:** 10.1101/2021.06.08.447450

**Authors:** Shuji Shinohara, Hiroshi Okamoto, Nobuhito Manome, Pegio-Yukio Gunji, Yoshihiro Nakajima, Toru Moriyama, Ung-il Chung

## Abstract

It has been stated that in human migratory behavior, the step length series may have temporal correlation and that there is some relationship between this time dependency and the fact that the frequency distribution of step length follows a power-law distribution. Furthermore, the frequency of occurrence of the step length in some large marine organisms has been found to switch between power-law and exponential distributions, depending on the difficulty of prey acquisition. However, to date it has not been clarified how the aforementioned three phenomena arise: the positive correlation created in the step length series, the relation between the positive correlation of the step length series and the form of an individual’s step length distribution, and the switching between power-law and exponential distributions depending on the abundance of prey. This study simulated foraging behavior using the Bayesian decision-making agent simultaneously performing both knowledge learning and knowledge-based inference to analyze how the aforementioned three phenomena arise. In the agent with learning and inference, past experiences were stored as hypotheses (knowledge) and they were used in current foraging behavior; at the same time, the hypothesis continued to be updated based on new experiences. The simulation results show that the agent with both learning and inference has a mechanism that simultaneously causes all the phenomena.

## 1. Introduction

Lévy walks are found in the migratory behavior of organisms at various levels, from bacteria and T cells to humans [1–6]. The Lévy walk is a random walk where the frequency of occurrence of a linear step length *l* follows a power-law distribution *P*(*l*) ~ *l^-η^*, 1 < *η* ≤ 3. Compared with the Brownian walk, which is also a random walk (the frequency of occurrence of the step length *l* is characterized by an exponential distribution (EP) *P*(*Z*) ~ *e^-λl^*), the Lévy walk is characterized by the appearance of linear movements over extremely long distances.

The reason behind the formation of such patterns in the movement of organisms has been investigated [7], and it has been found that they can be explained by the Lévy-flight foraging hypothesis [8,9]. This hypothesis states that when prey is sparse and randomly distributed and predators have no information (memory) about the prey, the Lévy walk becomes the optimal foraging behavior, whereas Brownian motion is theoretically optimal when prey is abundant [10,11]. In fact, it has been indicated that in a few marine organisms, when prey is abundant, a Brownian walk is observed, whereas when prey is scarce, a Lévy walk is observed [4,12]. Mechanisms for switching between Lévy and Brownian behavior have been the focus of considerable attention [13].

Humans have advanced learning and estimation capabilities and can acquire useful information through systematic search even in dynamic and uncertain nonstationary environments [14]. Ross et al. [15] demonstrated that in human exploration behavior, the mode of exploration changes depending on the encounter with prey. Particularly, they indicated that in response to encounters, hunters produce a more tortuous search of patches of higher prey density and spend a large part of their search time in patches; however, they adopt more efficient unidirectional, inter-patch movements after failing to encounter prey over a sufficiently long period. This type of search behavior is called area-restricted search (ARS) [15,16].

Wang et al. [17], Rhee et al. [18], and Zhao et al. [19] demonstrated that the temporal variation of step length is autocorrelated; this means that there is a trend in the time variation of step length, such that short (long) steps are followed by short (long) steps. Such a behavioral pattern, even if the frequency of occurrence of step length follows a power-law distribution, is not a pure random walk and therefore may not be called a Lévy flight. In addition, it has been emphasized that the dependence of the step length on time is related to the frequency distribution of the step length following a power-law distribution [17]. It is necessary to clarify the mechanisms that create these relationships.

Perhaps, in organisms with advanced cognitive functions, such as humans, various movement patterns may appear as a result of learning and inference through interaction with the environment, rather than as a result of random search [14,20]. In this study, we simulated the foraging behavior of predators based on the following two assumptions: 1) prey exist in two-dimensional space according to a density distribution like a normal distribution, and 2) predators have the ability to learn and infer the prey density distribution based on their own experience in acquiring prey.

Consider a situation where a predator estimates the distribution of prey using Bayesian inference—a machine learning method. In Bayesian inference, several candidate distributions are first prepared as hypotheses about how the prey are distributed. Next, confidence in each hypothesis based on observed data (experience of where and how many of the prey are actually acquired) is updated. It would be quite possible for predators to memorize locations where they have caught much food in the past as candidates (hypotheses) for foraging locations and use them in current and future foraging behavior. However, it is not possible to prepare an appropriate hypothesis at the initial stage when there is no information on prey. In addition, if the prey is an animal, it may move around, so there is no guarantee that the prey will be in the same place in the future. Furthermore, the consumption of food by predators could deplete the food supply in the area. In such a case, the hypothesis itself needs to be reformulated or revised based on observed data.

Gunji et al. [21–23] proposed a novel mathematical model called Bayesian and inverse Bayesian (BIB) inference that forms hypotheses based on observed data and subsequently performs inference using the hypotheses and observed data; in other words, learning and inference are performed simultaneously and seamlessly. The process of simultaneous learning and inference, similar to that in BIB inference, has been suggested in human-related experiments. Zaki and Nosofsky conducted a behavioral experiment that demonstrated the influence of unlabeled test data on learning [24]. Their experiment demonstrated that humans do not fix their hypothesis after training; instead, unlabeled test data influence the learned hypotheses of humans [25].

In this study, we propose a novel BIB inference model based on Bayesian inference with discount rate [26,27] as an algorithm for inference and the expectation-maximization (EM) algorithm [28,29], an unsupervised clustering method, as an algorithm for learning (inverse Bayesian inference). The model is used as a decision-making model for foraging agents. On the one hand, Bayesian inference is an algorithm that identifies the hypothesis that best explains the observed data, from one of several hypotheses (generating distributions) prepared in advance. On the other hand, the EM algorithm is an algorithm that, by modifying the initial distribution based on the observed data, approaches the distribution that best explains the observed data. Bayesian inference and the EM algorithm have the same goal of estimating the most appropriate generative model based on observed data. However, the direction of the estimation is opposite. If the estimated generative model is considered as knowledge, then the EM algorithm can be assumed to be a type of learning because it forms and modifies knowledge based on observed data. Further, Bayesian inference can be considered to be knowledge-based inference because it decides the best among the several hypotheses (generative models) prepared in advance.

In this study, we simulated the foraging behavior of BIB agents and analyzed the following phenomena: 1) the positive correlation in the step length time series, 2) the relationship between the positive correlation and the shape of the frequency distribution of the step length, and 3) the switching between the power law and the exponential distribution depending on the difficulty of food acquisition. The results of the foraging simulation showed that the mechanism of the BIB agent simultaneously causes the aforementioned phenomena.

## 2. Materials and Methods

### 2.1. Prey density distribution

In this study, we consider a situation wherein the prey are distributed in a two-dimensional space. We will deal with the case where prey are unevenly distributed in patches, and the case where prey are randomly distributed with uniform probability throughout the space (Figure 1). As an example of the former, we may consider the case where the prey is a plant or an animal with a nest. The plants can be unevenly distributed depending on the amount of sunlight in the area, the nutrients in the land, and the ground conditions (soil or rock). Furthermore, considering that animals roam around the nest, the closer to the nest, the higher the probability that animals are present. For the purpose of simplicity, we assume that the density (probability of existence) of prey in each patch is normally distributed. In other words, the prey density distribution of the patch *p* at a certain point *d* = (*d_x_*, *d_y_*) at time *t* is defined as follows:

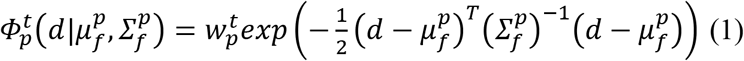

where 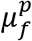 and 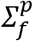 are the center and variance of patch *p*, respectively. 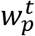 is a parameter that controls the total amount of prey of patch *p*, at time *t*. The distribution takes a maximum value when the condition 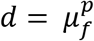 is satisfied, and the value is 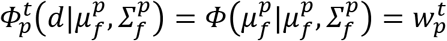. Moving away from the center 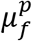 of the distribution, the prey density of decreases. 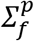 is a variance parameter, and the larger 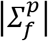 is, the more widely the prey are distributed and the larger is the total number of prey.

**Figure 1.**
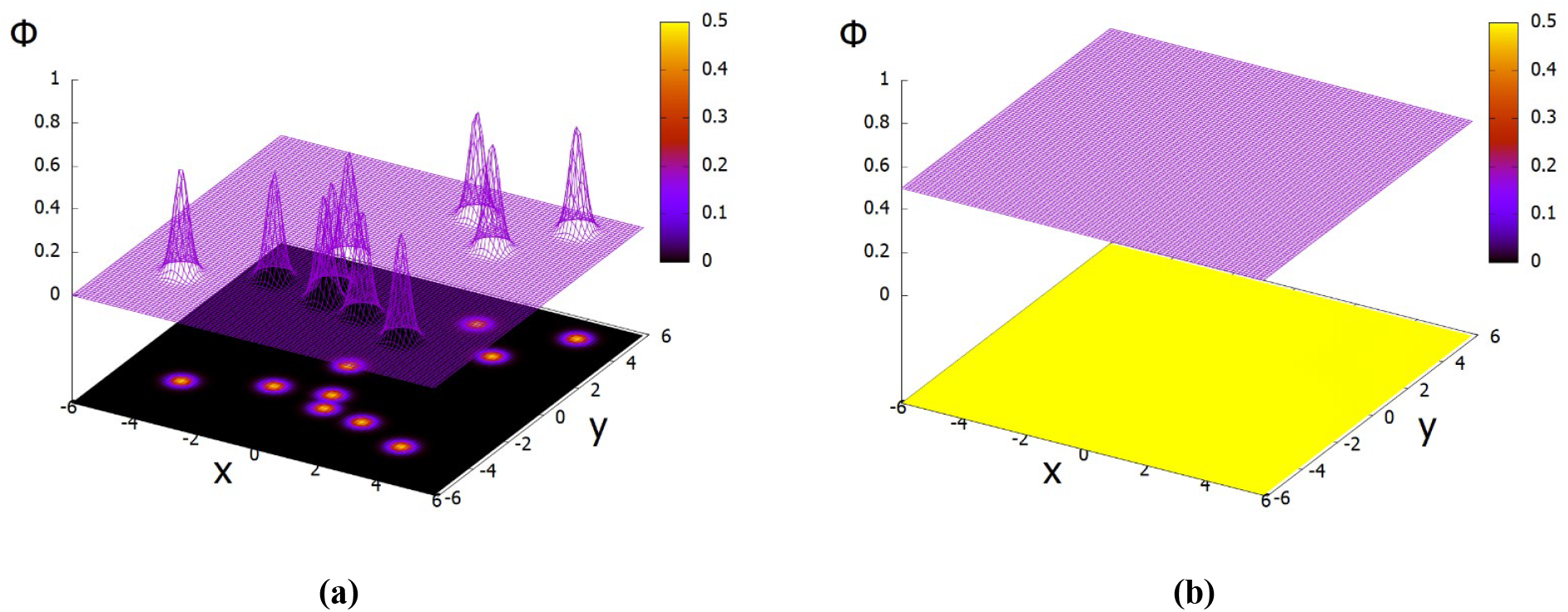
Examples of prey density distribution. Predator probabilistically acquires prey in proportion to the prey density at the foraging point. (a) Distribution in the case of 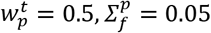. (b) Distribution in the case of 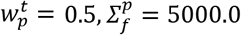. This represents an approximate uniform distribution.

### 2.2. Predator agent with Bayesian and inverse Bayesian (BIB) inference

#### 2.2.1 Overview

The main objective of the agent is to acquire as many preys as possible within a certain period. The agent has the ability to learn and estimate the prey density distribution in order to acquire food efficiently. The agent has several candidate of prey density distributions as hypotheses, and the correct hypotheses are estimated based on the food acquisition information. In other words, the agent updates its confidence in each hypothesis. At the same time, the agent modifies the contents of each hypothesis, that is, the parameters (model) of the prey density distribution corresponding to each hypothesis. The Bayesian inference is used to update the confidence level. The EM algorithm is used to modify the hypothesis models. These are described in detail in the next subsection.

The agent decides probabilistically at which point 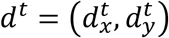 to forage according to the model of the hypothesis 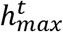, with the highest confidence at time *t*, as follows:

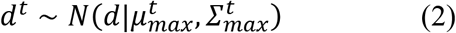

where 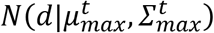 is the model of the hypothesis wherein the agent is most confident, and is represented by a normal distribution with 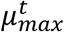 as the centre and 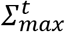 as the variance. Next, whether a prey is acquired is probabilistically determined based on the prey density at point *d^t^*.

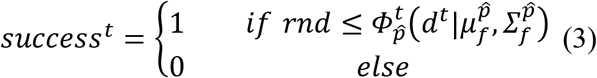

where *rnd* represents a uniform random number in the [0,1] interval; *success^t^* = 1 represents successful prey acquisition, and *success^t^* = 0 represents unsuccessful prey acquisition. The foraging patch 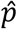 is defined as the patch with the highest prey density at the foraging site *d^t^*. In other words,

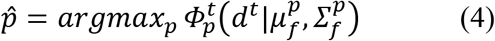

It is assumed that the agent consumes food every time it acquires, and the prey amount is set to decrease by Δ. If the agent acquires prey at the location *d^t^* at time *t*, then 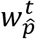 in the patch 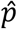 decreases as follows:

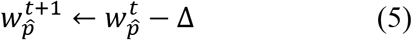

The following points should be noted here. *Φ,* defined in Equation (1), is the actual prey density distribution, which varies depending on the consumption of food. On the other hand, the *N* used in Equation (2) is the prey distribution estimated by the agent. Therefore, this *N* changes according to the agent’s learning and estimation based on the information of food acquisition.

#### 2.2.2. Sequential discounting expectation-maximization (SDEM)

In this subsection, we describe how to modify the hypothetical model. The EM algorithm is used to modify the model. The EM algorithm [28] is a machine learning technique used for unsupervised clustering. The classification of a given set of *n* data {*d*^1^, *d*^2^, …, *d^n^*} into *M* clusters is considered here. First, a mixture normal distribution comprising *M* normal distributions is considered for generating data *d*.

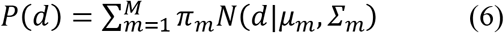

where d is a D-dimensional data vector and *N*(*d|μ_m_,∑_m_*) is a D-dimensional normal distribution with mean *μ_m_* and variance-covariance matrix *∑_m_*, as shown below.

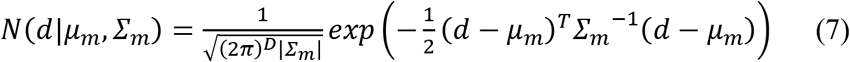

Here, 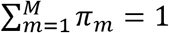 is assumed. *π_m_* is called the “mixing weights” and represents the weight of each normal distribution. The aim of the EM algorithm is to estimate the mixing weights *π_m_*, mean *μ_m_*, and variance *∑_m_* of *M* normal distributions from *n* data {*d*^1^, *d*^2^, …, *d^n^*}. First, in Step E, the responsibility is calculated. The responsibility for the normal distribution *m* of data *d^i^* is calculated as follows:

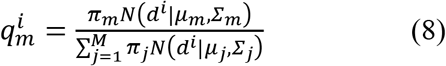

Next, the weights of data *d^i^* with respect to the normal distribution *m* are defined as follows:

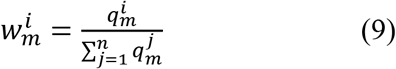

Next, in Step M, the mixing weights, mean, and variance of each normal distribution are updated.

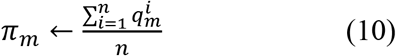

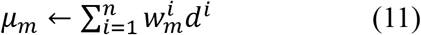

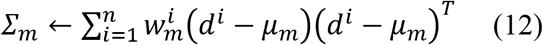

In the EM algorithm, the values converge by alternately repeating Steps E and M. However, in the EM algorithm, it is necessary to provide all observational data simultaneously. In practice, there are cases where data processing must be performed sequentially, each time after the data are observed. Various online algorithms have been proposed to address this situation [30–37]. For example, Yamanishi et al. [29] proposed an SDEM algorithm. In the SDEM algorithm, Steps E and M are executed once for each data observed sequentially. In addition, the SDEM algorithm introduces the effect of forgetting to deal with unsteady situations where the inferred target changes. Specifically, weighting is performed to weaken the influence of the older observation data by introducing a discount rate β (0<β<1). When data *d^t^* are observed at time t, the mixing weights, mean, and variance of each normal distribution are updated, as shown below.

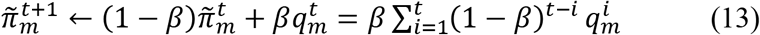

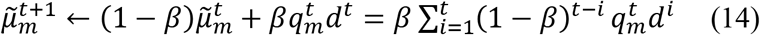

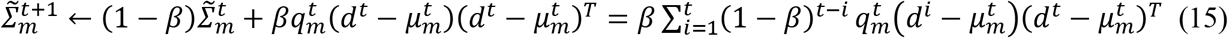

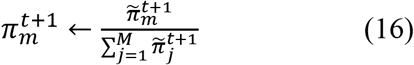

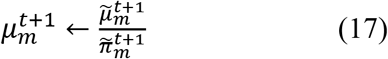

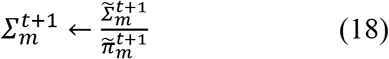

In this study, for simplicity, the case of a single normal distribution, that is, *M*=1, is treated as the generating distribution of the data. In the case of *M*=1, 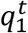 and 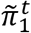 are always one. For simplicity, 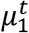 and 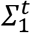 are denoted as *μ^t^* and *∑^t^*, respectively. At this time, Equations (17) and (18) show that the new estimated value is obtained as a convex combination of the current estimated value and current observed data, as follows:

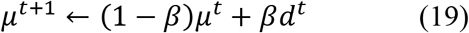

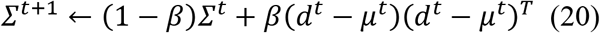

Equations (19) and (20) represent the exponential moving averages of *μ^f^* and *∑^t^*, respectively.

#### 2.2.3. Bayesian inference with discount rate (BID)

In this subsection, we describe how to update the confidence level for each hypothesis. The Bayesian inference is used to update the confidence level. In Bayesian inference, first, several hypotheses (generative models) are prepared for an inferred target. Next, the evidence (the data generated by the target) is observed and the confidence in each hypothesis is updated by quantitatively evaluating the fit between each hypothesis and the observed data. Finally, based on the confidence level, the single best hypothesis is determined.

In general, it is better to obtain as much observational data (information) as possible for the accurate estimation of the target. However, this is true only in a stationary environment. When the target varies dynamically, it is necessary to discard the distant past observation data for a more accurate estimation. To address this situation, BID is introduced here [26,27].

First, a discrete version of Bayesian inference is described. In Bayesian inference, first a number of hypotheses *h_k_* are defined and a model (the generating distribution of the data) for each hypothesis is prepared in the form of a conditional probability *P*(*d|h_k_*). Specifically, d is assumed to be a D-dimensional vector, as in the case of SDEM, and the model *P*(*d|h_k_*) is assumed to be a D-dimensional normal distribution, as follows:

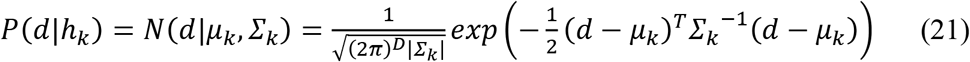

This conditional probability is called the likelihood if the data are fixed and considered as a function of the hypothesis *h_k_*; that is, the parameter *θ_k_* = (*μ_k_, ∑_k_*) of the normal distribution. An initial confidence *P^1^*(*h_k_*) value for each hypothesis is provided as a prior probability. If the confidence in hypothesis *h_k_* at time t is *p^t^*(*h_k_*) and data *d^t^* are observed, the posterior probability *P^t^*(*h_k_|^l^*) is calculated using Bayes’ theorem, as follows:

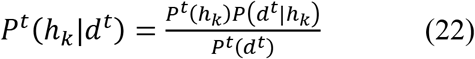

where *p^t^* (*d^t^*) is the marginal probability of the data at time t and is defined as follows:

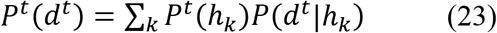

The following Bayesian update accordingly replaces the posterior probability with the confidence the next time.

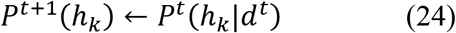

Combining Equations (22) and (24), we obtain Equation (25):

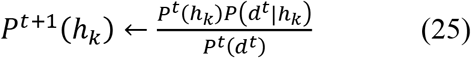

The estimation proceeds by updating the confidence in each hypothesis by Equation (25), each time the data are observed. Here, the confidence in each hypothesis satisfies *∑_k_ P*^*t*+1^(*h_k_*) = 1. Focusing on the recursive nature of *P^t^*(*h_k_*), Equation (25) can be rewritten as follows:

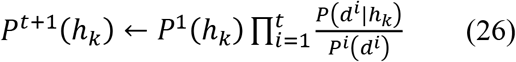

Here, *P^i^*(*d^i^*) is common to all hypotheses and can be regarded as a constant. Omitting the normalization process, Equation (26) can be written as follows:

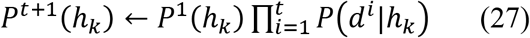

That is, the current confidence of each hypothesis is proportional to the prior probability multiplied by the likelihood of the data observed each time.

Next, the forgetting function to deal with unsteady situations is incorporated by introducing a discount factor into the Bayesian inference, as follows [26,27]. However, in the following, *C* is used instead of *P*, to distinguish between Bayesian inference and BIB inference, by introducing a discount factor. If the normalization process is omitted, the updating formula for the confidence of each hypothesis is defined as follows:

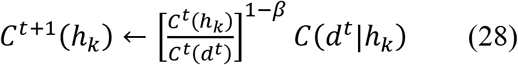

where *β* is the discount factor and Equation (28) agrees with Equation (25) when *β* = 0. After Equation (28), normalization is conducted to satisfy *∑_k_ C*^*t*+1^(*h_k_*) = 1, as follows:

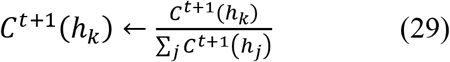

If we substitute *t*-1 for *t* in equation (28), we get

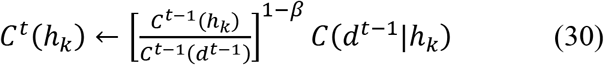

Therefore, Equation (28) can be transformed using Equation (30) as follows:

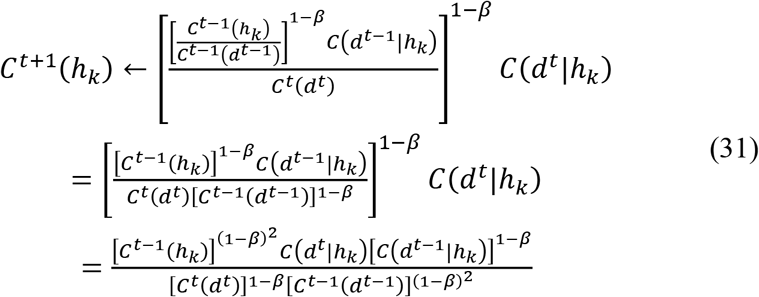

By repeating the above operation recursively, we finally obtain the following equation.

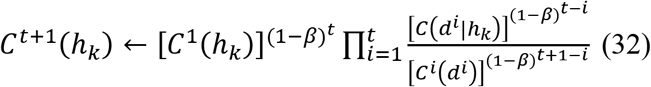

In Equation (32), the denominator *C^i^*(*d^i^*) on the right-hand side is common to each hypothesis and can be regarded as a constant; therefore, omitting the normalization process, the following equation is obtained:

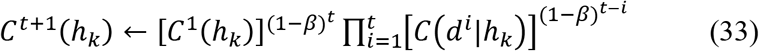

In other words, it is understood that the current confidence is multiplied by the likelihood in the distant past with less weight. When *β* = 0, using *C*^*t*+1^(*h_k_*) ← *C*^1^(*h_k_*)*C*(*d*^1^|*h_k_*)*C*(*d*^2^|*h_k_*) – *C*(*d*^1^ |*h_k_*) in the Bayesian inference, the current likelihood and the likelihood in the distant past are evaluated with equal weight. When *β* = 1, *C*^*t*+1^(*h_k_*) ← *C*(*d*^1^|*h_k_*) is used to calculate the next confidence using only the likelihood of the current observation. In the BID, the confidence for each hypothesis is updated according to Equation (28), each time data are observed, and the best hypothesis 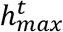 is determined based on the confidence level.

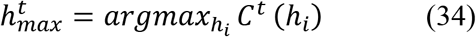

#### 2.2.4. BIB inference

SDEM and BID are already explained. If the generative model is regarded as knowledge, SDEM can be regarded as a type of learning because knowledge is formed and modified based on the observed data. BID, however, can be called knowledge-based inference because it decides which of the several hypotheses (generative models) prepared in advance is the best.

In this section, a model of BIB inference is proposed that employs SDEM and BID to perform learning and inference simultaneously. Here, BID and SDEM are the models for Bayesian inference and inverse Bayesian inference, respectively. This is a model where the hypotheses are revised based on inverse Bayesian inference, while Bayesian inference is used to assess which hypothesis is correct. Because the model *C*(*d|h_k_*) of the hypothesis *h_k_* changes with time through inverse Bayesian inference, it is denoted by (*d|h_k_*), with time t as a subscript.

It is assumed that food is obtained by foraging at point *d,t* at time t. At this time, the confidence of each hypothesis is updated using the BID equation:

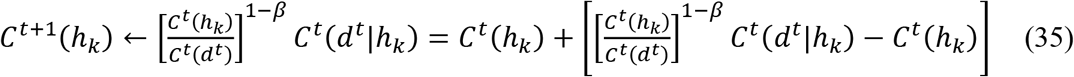

On the other hand, when it fails to acquire food at *d^t^*, the confidence level is updated as follows:

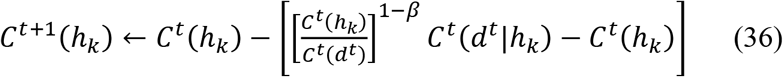

In other words, contrary to the successful case, the higher the likelihood of the hypothesis for *d^t^*, the lower the confidence level in the hypothesis.

When the food is successfully acquired, the hypothesis 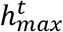 with the highest confidence is selected using Equation (34). SDEM is thereafter used to update the model 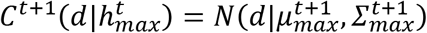, for the following hypothesis 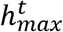, as follows.

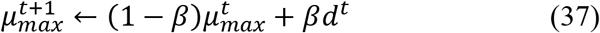

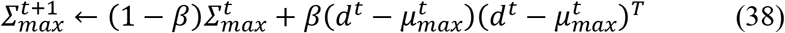

This is repeated every time data are observed. The group of processes described in this section is summarized as the following algorithm:

1. Determine the parameters 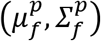 of the prey density distribution.
2. Set the values for parameters *β, K*.
3. Establish the initial values for 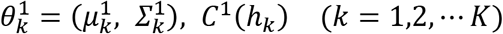.
4. Repeat the following.

4.1 The agent chooses the hypothesis 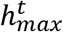 it believes in the most, according to Equation (34).
4.2 Based on the model of the hypothesis 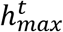, the agent decides the foraging point according to Equation (2).
4.3 Equation (3) determines the success or failure of foraging.
4.4 The agent executes the following processes simultaneously:

4.4.1 Bayesian inference (BID): The confidence *C^t^*(*h_k_*) of each hypothesis is updated according to Equation (35) or (36).
4.4.2 Inverse Bayesian inference (SDEM): The model 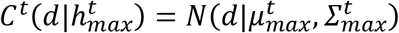 of the hypothetical 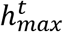 is modified according to Equations (37) and (38) only if the prey is successfully acquired.

### 2.3. Parameter setting

The dimensionality of the data was set to *D*=2 because we are simulating foraging behavior in a two-dimensional space. The number of hypotheses was set as *K*=15. For the initial values of the model parameters for each hypothesis, the mean 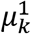 was determined randomly in the range [-5,5] for both the x and y coordinates. The variance was set as 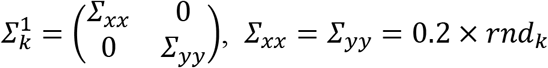, where *rnd_k_* represents a uniform random number in the [0,1] interval. The initial value of confidence (that is, prior probability) for each hypothesis was set as *C*^1^(*h_k_*) = 1/*K*, such that they would be equal. For all agents, the discount rate was set as *β* = 0.03. We set the target two-dimensional space in the range [-5,5] for both x and y coordinates. If a larger range is set, a larger number of hypotheses *K* and the variance 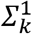 need to be set in order to cover the entire space. We also set the discount factor *β* = 0.03. If the food consumption rate *Δ* becomes large, the prey density distribution will fluctuate greatly, and thus, it is necessary to set a larger discount for efficient foraging behavior.

For prey density distributions, five cases were simulated: from scarce prey to abundant prey. First, the central coordinates 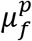 of prey distribution were randomly determined in the range [-5,5] for both the x and y coordinates. The variances of the prey distribution were set as 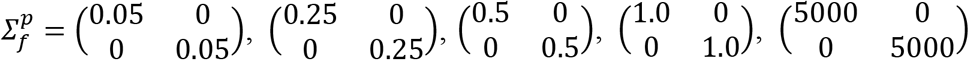. A distribution with 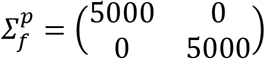 represents an approximate uniform distribution. 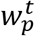 is a parameter that controls the total amount of prey of patch *p*, at time *t*. Here, 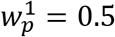 was set as the initial value. The number of patches was set to 10. The food consumption rate Δ was set to 0.0005. The simulation was conducted up to the 30000^th^ step.

The simulation program was developed based on the C++ language. The compiler was MinGW 8.1.0 64-bit for C++ [38]. The Qt library (Qt version Qt 5.15.2 MinGW 64-bit) was also used for development [39].

### 2.4. Autocorrelation coefficient and correlation time

In this study, autocorrelation coefficient and correlation time are used to measure the time dependence of step-length time series. For a given *T* time-series data (*l*_1_, *l*_2_, – *l_T_*), the autocorrelation coefficient r(τ) with time lag τ is expressed as follows:

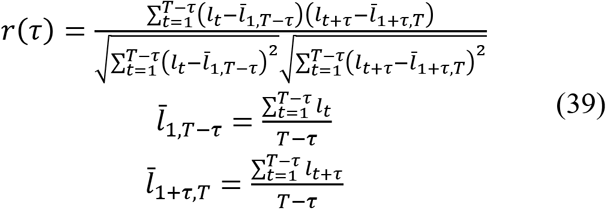

In this study, the autocorrelation coefficient was obtained in the range *τ* = 1-2500. The autocorrelation coefficients for every time-series data are averaged from *r*(1) to *r*(2500) and are expressed as 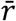.

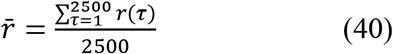

Thereafter, 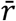 is referred to as correlation time.

### 2.5. Statistical analyses

The following analyses were performed using the statistical software R, version 3.6.1 (2019-07-05) [40], unless otherwise specified. We used the R packages of RVAideMemoire version 0.9.81 for the pairwise comparisons using Fisher’s exact test and multcomp version 1.4.16 for the Tukey–Kramer test. The operating system used was Windows 10 (Microsoft Corp., Redmond, WA, USA). For all analyses statistical significance was set at p < 0.05.

## 3. Results

We performed foraging simulations for five cases from scarce prey to abundant prey. The agent probabilistically decided where to forage according to its own estimated model that was represented by a normal distribution in this study. Prey were acquired stochastically in proportion to the prey density at the foraging point. The objective of the agent was to acquire as many prey as possible within a certain period.

Figure 2 shows an example of the foraging behavior of an agent in the case of 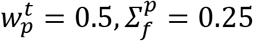. Figure 2(a) plots the foraging points at each time point. The z axis and color represent time steps. In the figure, two points *d*^*t*-1^ and *d^t^* at adjacent times are connected by straight lines. Figures 2(b) and 2(c) show examples of the prey distribution at the initial stage (*t* = 1) and *t* = 3000, respectively. It can be seen that some patches are on the verge of disappearing due to the consumption of food. Figure 2(d) shows the time series of patch numbers explored by the agent. Figure 2(e) shows the time evolution of the expected reward, expressed in terms of prey density 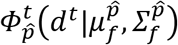 at the foraging point *d^t^*.

**Figure 2.**
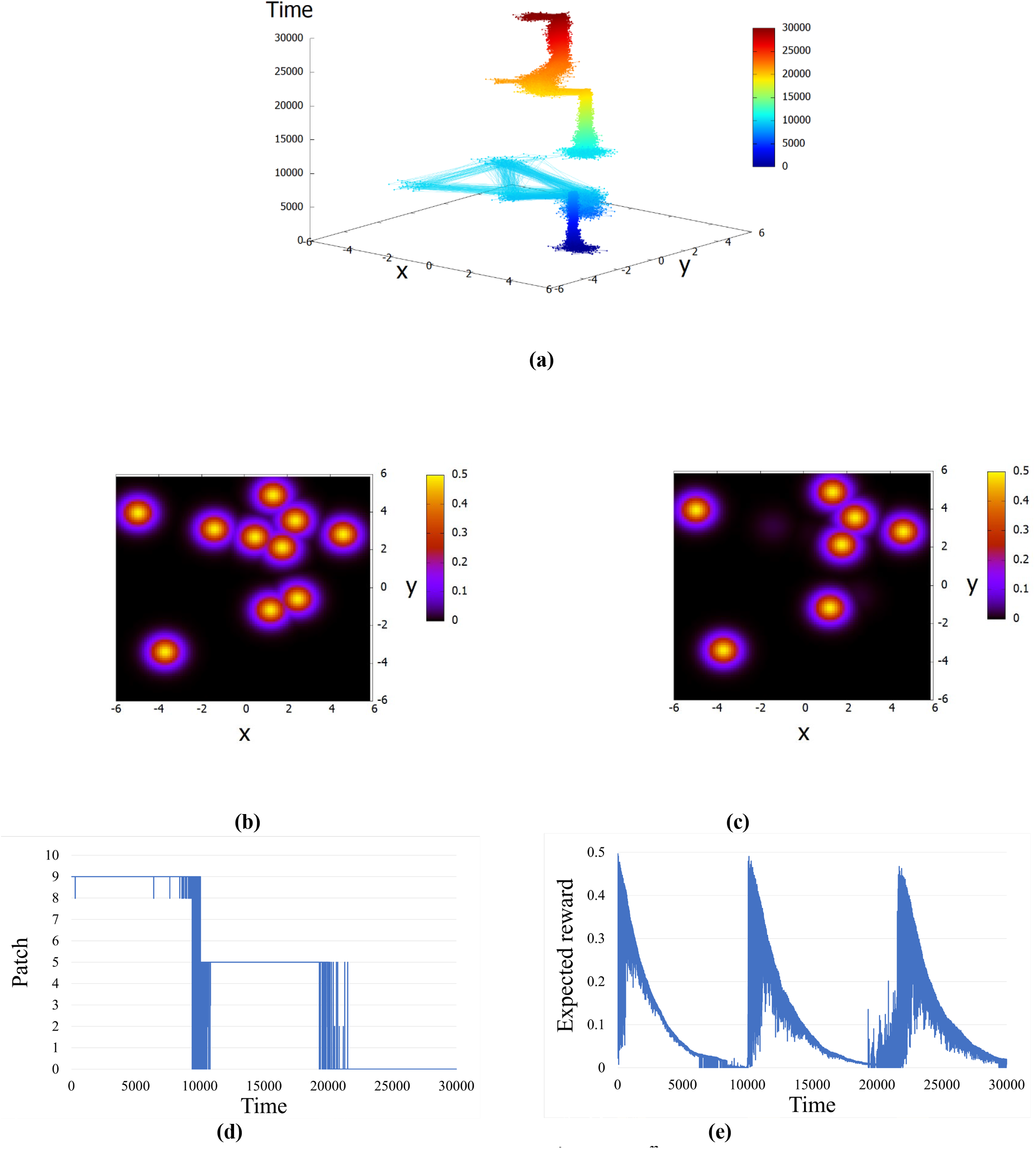
Example of foraging behavior in the case of 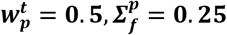. (a) Movement pattern from time 1 to 30000. Two foraging points at adjacent time points are connected by a straight line. Z axis and color represent time steps. (b) Prey density distribution at the initial stage (t = 1) (c) Prey density distribution at t = 30000. (d) Time evolution of patch numbers explored by an agent. (e) Time evolution of the expected reward.

As can be seen in these figures, when an agent finds a patch, it forages intensively in that patch. When the food is consumed and the amount of food in the patch becomes low, the agent repeats the process of searching and moving to a new patch.

Figures 3(a) and 3(b) show examples of the time evolution of the step length *l_t_* = ||*d_t_* – *d*_*t*-1_||_2_. Figures 3(c) and 3(d) show the complementary cumulative distribution function (CCDF) of the step length *l_t_* when Figures 3(a) and 3(b), respectively. These figures also show the results of fitting a truncated power-law distribution (TP) model (green) and an EP model (red) to the simulation data. Of the step length data, only the data within the range 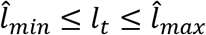 are used to improve the fitting as much as possible. See the Supplementary Information (SI) for details on how the fitting range 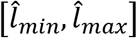 and fitting method are determined. The CCDF graph is such that the CCDF value when 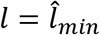, is set to one. Figures 3(b) and 3(d) are double logarithmic graphs with both axes displayed in logarithmic form.

**Figure 3.**
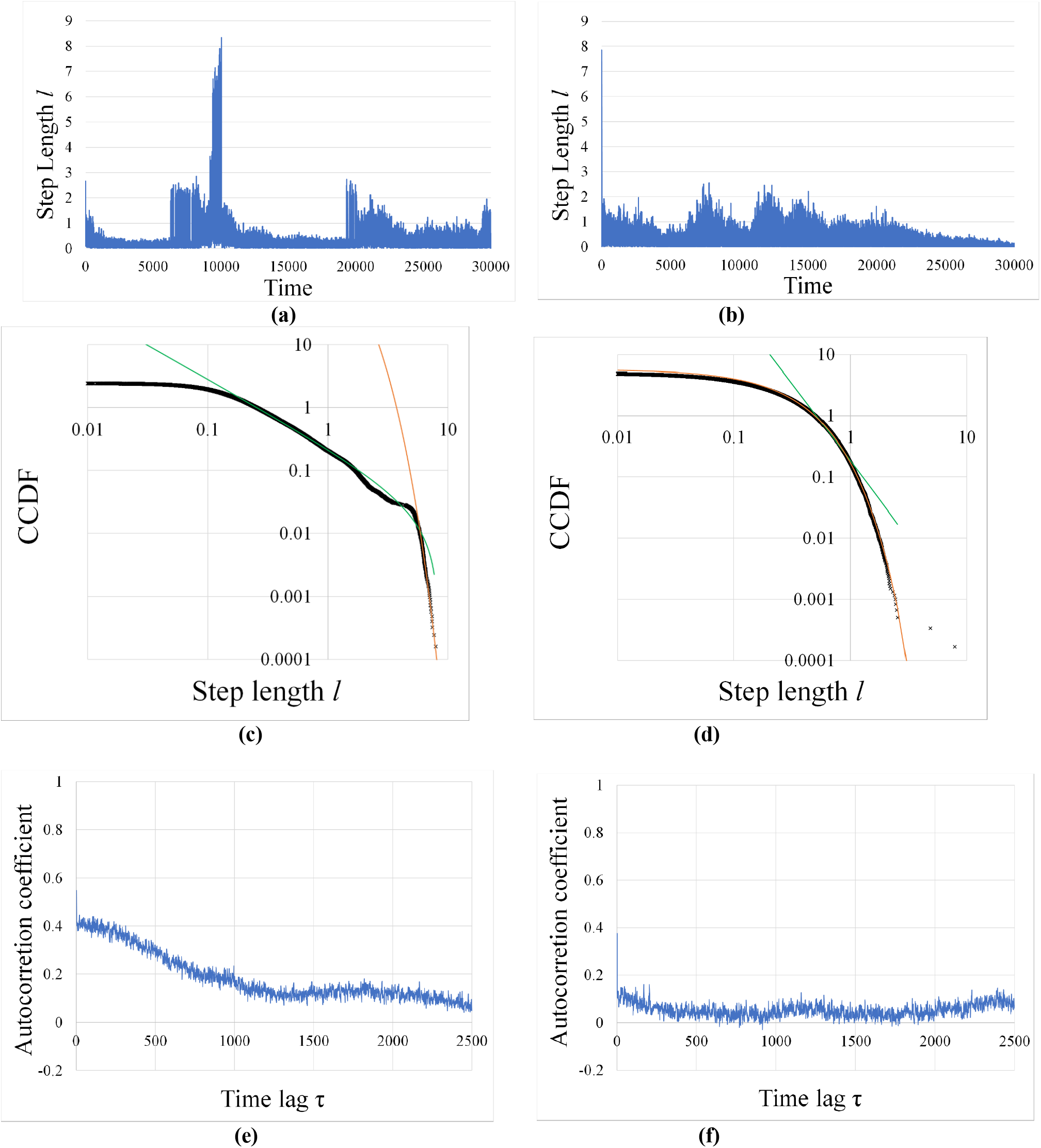
Examples of time evolution of step length, Complementary cumulative distribution function (CCDF) of step length, and autocorrelation of step length time series. (a) Time evolution of step length in the case where a Lévy walk-like gait pattern is observed. (b) Time evolution of step length in the case where a Brownian walk-like gait pattern is observed. (c) CCDF of the step length in the case of (a). This figure also shows the results of fitting a TP model (green) and an EP model (red) to the simulation data. Exponent of the TP is *η* = 2.08. Fitting range is [0.25, 8.34]. (d) CCDF of the step length in the case of (b). This figure also shows the results of fitting a TP model (green) and an EP model (red) to the simulation data. Exponent of the EP is *λ* = 3.53. Fitting range is [050, 7.84]. (e) Correlogram for the step-length time series shown in (a). (f) Correlogram for the step-length time series shown in (b).

As seen from these figures, a Lévy walk-like gait pattern, wherein the frequency distribution of the step length shows a power distribution, and a Brownian walk-like gait pattern, which shows an exponential distribution, appear. Figures 3(e) and 3(f) show the correlogram of the step-length time series shown in Figures 3(a) and 3(b), respectively. The horizontal and vertical axes represent the time lag τ and the autocorrelation coefficient for each *τ*, respectively. From these figures, we can see that the temporal autocorrelation is stronger for Lévy walk-like gait pattern.

Hereafter, we will refer to the group of agents whose frequency distribution of step length is classified as EP as the EP group, and the group of agents whose frequency distribution of step length is classified as TP as the TP group. Furthermore, we will divide the TP group into those whose exponent *η* is in the range 1.0 ≤ *η* ≤ 3.0 (TP1) like the Levy walk and those whose exponent *η* is not in the range of 1.0 ≤ *η* ≤ 3.0 (TP2). Table 1 shows the number of times the frequency distribution of the step length has been classified as TP1, TP2, and EP, as the results of 100 simulation trials with different random seeds for each of the cases of the five prey density distributions (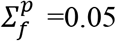, 0.25, 0.5, 1.0, 5000.0).

**Table 1.** Number of trials for which the frequency distribution of the step lengths was classified as TP1, TP2, and EP.

| Group | Variance of prey density distribution |  |  |  |  |
| --- | --- | --- | --- | --- | --- |
| | $\Sigma_f^p = 0.05$ | $\Sigma_f^p = 0.25$ | $\Sigma_f^p = 0.5$ | $\Sigma_f^p = 1.0$ | $\Sigma_f^p = 5000.0$ |
| TP1 | 62 | 62 | 30 | 4 | 7 |
| TP2 | 12 | 14 | 15 | 9 | 2 |
| EP | 26 | 24 | 55 | 87 | 91 |
| Total | 100 | 100 | 100 | 100 | 100 |

Fisher’s exact test with Bonferroni correction was performed for the ratio of TP1 to EP. The results showed that there were significant differences in the ratio at 0.1% level for all combinations except 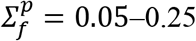 and 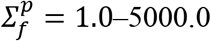. In other words, the scarcity of food tends to increase the percentage of TP1 compared to EP.

Figures 4 shows the time dependence of the step length. Figure 4(a) shows the mean values of correlation time for each group shown in Table 1 for each prey density distribution. Two-way ANOVA of correlation time was conducted for two factors, groups (i.e., EP, TP1, or TP2) and variances of prey density distribution (i.e., 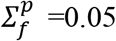, 0.25, 0.5, 1.0, 5000.0). Significant differences were noted in both group and variance factors (F(2, 485) = 8.83, p < 0.001, F(4,485) = 73.16, p < 0.001). However, there was no interaction between them (F (8, 485) = 1.50, p = 0.154).

**Figure 4.**
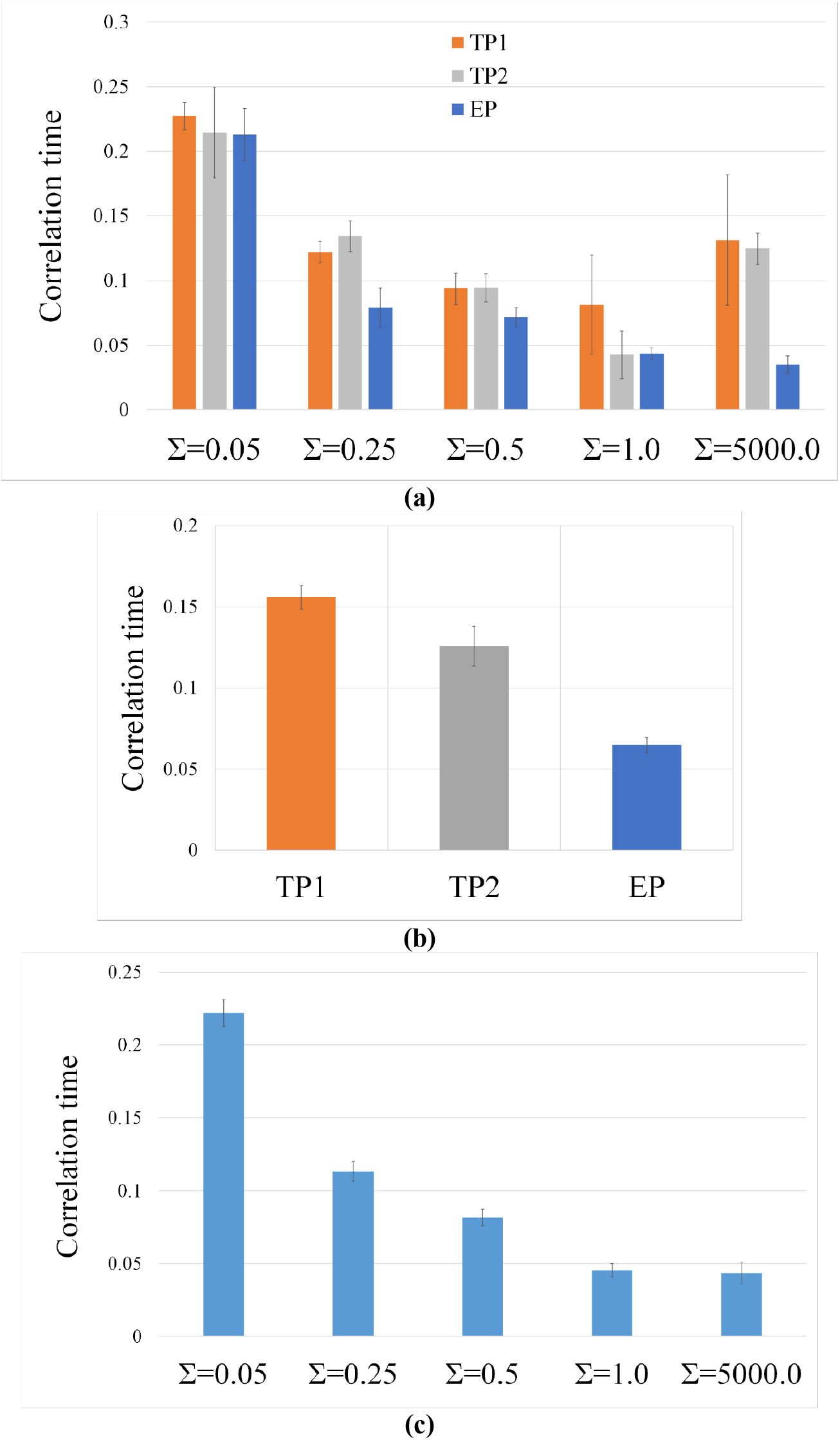
Comparison of correlation time. (a) Average correlation time for each group shown in Table 1 for each prey density distribution. (b) Average correlation time for each group. (c) Average correlation time for each prey density distribution. In all figures, the error bars represent the standard error.

Figure 4(b) shows the average correlation time for each group. The Tukey–Kramer test revealed significant differences between the EP and the TP1 groups (p < 0.001) and between the EP and the TP2 groups (p < 0.001). However, there was no significant difference between TP1 and TP2 groups (p = 0.070). The correlation time of TP1 is larger than that of EP, i.e., there is a stronger time dependence in the time variation of the step length for the agents belonged in TP1 group.

Figure 4(c) shows the average correlation time for each prey density distribution. The results of the Tukey-Kramer test showed that there are significant differences in correlation time among all variances except between 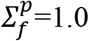 and 5000.0 at the level of 5%. Thus, the more abundant the food, the smaller the correlation time. Conversely, the sparser the food, the stronger the time dependence of the step length.

## 4. Discussion

In this study, a novel model of BIB inference was proposed wherein knowledge learning and knowledge-based inference are performed simultaneously. The learning was modeled using SDEM, an online EM algorithm, and the inference was modeled using BID, a Bayesian inference with discount factor. Next, foraging simulations were conducted using a decision-making agent incorporating the BIB inference model. In BIB agents, past experiences are stored as hypotheses (knowledge) and are used in the current foraging behavior; at the same time, the hypothesis continues to be updated based on new experiences.

The following are the three results of the simulations. First, when prey are sparse, the frequency distribution of the step length (that is, the distance between the foraging point at one time and the foraging point the next time) tends to follow a TP, whereas when prey are dense, the distribution tends to become an EP. This result is consistent with the finding that the frequency of occurrence of the step length in some marine organisms switch between power-law and exponential distributions depending on the amount of prey [4,12]. Second, the time variation of the step length tends to show a strong time dependence when the frequency distribution of the step length follows a power-law distribution. This result is consistent with the finding [17] that there is a relationship between the dependence of the step length on time and the fact that the frequency distribution of the step length follows a power-law distribution. Third, the more sparsely distributed the prey, the stronger the time dependence of the step length.

When the frequency distribution of the step length follows a TP, the search behavior of BIB agent is divided into two modes: a rough search to select the patch and a detailed search to detect the center of the selected patch. This switch may be considered a model of ARS. The agents show extremely long steps in the patch search mode. However, after fixing the patch, gradually, the agent will choose a location near the center of the patch as its foraging point in the detailed search mode. Accordingly, the step length approaches zero. Because the step length tends to decrease gradually, there is a stronger time dependence of the step length in the TP group. It is interesting to note here that although the foraging points are sampled from a normal distribution as shown in Equation (2), the frequency distribution of the distances between foraging points, that is, the step lengths, results in a TP.

In the case of prey abundance, the frequency distribution of the step length tends to become an EP. In the case of prey abundance, it is not necessary to cause the center of the estimated model to coincide with the center of the prey distribution and to converge the variance of the estimated model to zero, because prey can be obtained with a high probability without foraging at the center of the patch. Because the foraging point does not gradually move closer to the patch center, the time dependence is expected to be weaker with respect to the temporal variation of the step length. Thus, it can be noted that depending on the difficulty of prey acquisition, various migratory behavioral patterns emerge.

In general, the EM algorithm or Bayesian inference attempts to accurately estimate the parameters (mean and variance) of the generating distribution that produces the observed data. It is important to note that the observed data are usually sampled from the generating distribution; therefore, regardless of how the EM algorithm or Bayesian inference estimates the generating distribution, it has no effect on the observed data. However, in the simulation task of this study, foraging points are determined based on the model estimated by the agent; this implies that the observed data depend on the distribution that the agent estimates. This can lead to the variance of the model estimated by the BIB agent approaching zero, even though the variance of the actual prey distribution is not zero.

The exploration and preferential return (EPR) model incorporates the random walk process of exploration and the human propensity to revisit places (preferential return). In EPR, one of these two competing mechanisms is chosen probabilistically each time [41]. Exploration and exploitation are two essential components of any optimization algorithm [42]. Determining an appropriate balance between exploration and exploitation is the most challenging task in the development of any metaheuristic algorithm, owing to the randomness of the optimization process [43]. In the present simulations, two patterns emerged: a rough search using knowledge (hypotheses) and a detailed search where the target was narrowed down by modifying the knowledge. The former may be associated with exploitation and the latter with exploration. It is interesting to note that the emergence of these behavioral patterns in BIB agents is not explicitly controlled by any parameters. As can be seen from the simulation procedure, the BIB agent performs both learning and inference simultaneously at every step.

The limitation of this study is that the simulations addressed only the case of a single cluster with respect to the mixture distribution of the SDEM. In future work, we intend to perform simulations using a generative distribution comprising several normal distributions. In the present study, we regarded the generation distribution and density distribution of prey in each patch as a normal distribution. In future work, we intend to simulate a case wherein the density distribution of the prey inside the patch is uniform or nonparametric.

## Funding

This work was supported by the Center of Innovation Program from the Japan Science and Technology Agency, JST; and JSPS KAKENHI [grant number JP21K12009].

## Competing interests

N.M. is employed by SoftBank Robotics Group Corp. This does not alter our adherence to *Chaos, Solitons & Fractals* policies on sharing data and materials. All other authors declare no conflicts of interest.

## Data availability statement

The source code and data used to produce the results and analyses presented in this manuscript are available from Bitbucket Git repository: https://bitbucket.org/ShujiShinohara/levyflightforagingsimulation1/src/master/

## Notes

### Competing Interest Statement

The author [N. M.] is employed by SoftBank Robotics Group Corp. This does not alter our adherence to bioRxiv policies on sharing data and materials. All other authors declare no conflicts of interest.

