## Supplementary material for "Simulation of foraging behavior using a decision-making agent with Bayesian and inverse Bayesian inference: Temporal correlations and power laws in displacement patterns": SI

#### Fitting to simulation data

Here, a method is described to fit the frequency distribution of the data observed by simulation to the truncated power-law distribution (TP) model and the exponential distribution (EP) model. Here, the case in which the data take continuous values is discussed. For discrete values, please refer to [1].

##### Fitting to TP

Here, a method is described to fit the frequency distribution of the step length $l$ observed by simulation to the TP. The method is based on previous studies [2–6]. Specifically, the aim is to determine the minimum $\hat{l}_{\min}$ and maximum values $\hat{l}_{\max}$ of the observed data to be fitted to the TP model and the exponent $\hat{\eta}$ of the TP model that best fits the data in the range $\hat{l}_{\min} \le l \le \hat{l}_{\max}$. First, $\hat{l}_{\max}$ is the longest step length of the observation data. Next, the calculation method $\hat{l}_{\min}$ is described. In the case of a continuous distribution, TP in the range $l_{\min} \le l \le l_{\max}$ is expressed by the following formula:

$$p\left(l;\eta,l_{\min},l_{\max}\right)=\frac{\eta-1}{l_{\min}^{1-\eta}-l_{\max}^{1-\eta}}l^{-\eta} \qquad \text{(S1)}$$

The complementary cumulative distribution function (CCDF) of $p\left(l;\eta,l_{\min},l_{\max}\right)$ is expressed in the following equation.

$$P\left(l;\eta,l_{\min},l_{\max}\right)=1-\frac{l_{\min}^{1-\eta}-l^{1-\eta}}{l_{\min}^{1-\eta}-l_{\max}^{1-\eta}}=\frac{l^{1-\eta}+l_{\max}^{1-\eta}}{l_{\min}^{1-\eta}-l_{\max}^{1-\eta}} \quad \text{(S2)}$$

If the observed data in the range of $l_{\min} \le l \le l_{\max}$ are $\{l_1, l_2, \cdots, l_n\}$, then the log-likelihood of these data for TP is calculated using Equation (S1) as follows:

$$L\left(\eta;l_{\min},l_{\max}\right)=\sum_{i=1}^{n}\ln p\left(l_i;\eta,l_{\min},l_{\max}\right)=n\left(\ln\left(\eta-1\right)-\ln\left(l_{\min}^{1-\eta}-l_{\max}^{1-\eta}\right)\right)-\eta\sum_{i=1}^{n}\ln l_i \quad \text{(S3)}$$

The exponent $\hat{\eta}\left(l_{\min},l_{\max}\right)$ of the TP model that best fits the data in the range $l_{\min} \le l \le l_{\max}$ is $\eta$ that maximizes $L\left(\eta;l_{\min},l_{\max}\right)$. Specifically, $\eta$ is varied from 0.5 to 3.5, in increments of 0.01, to obtain $\hat{\eta}\left(l_{\min},l_{\max}\right)$ that numerically maximizes Equation (S3).

The authors introduce the Kolmogorov-Smirnov statistic $D\left(l_{\min},l_{\max}\right)$ to measure the closeness of CCDF $S\left(l;l_{\min},l_{\max}\right)$ obtained from the data in the range $l_{\min} \le l \le l_{\max}$ and the theoretical CCDF $P\left(l;\hat{\eta}\left(l_{\min},l_{\max}\right),l_{\min},l_{\max}\right)$ represented by Equation (S2).

$$D\left(l_{\min},l_{\max}\right)=\max_{l_{\min}\le l\le l_{\max}}\left|S\left(l;l_{\min},l_{\max}\right)-P\left(l;\hat{\eta}\left(l_{\min},l_{\max}\right),l_{\min},l_{\max}\right)\right| \quad \text{(S4)}$$

If $l_{\max}=\hat{l}_{\max}$ is fixed, then $D\left(l_{\min},\hat{l}_{\max}\right)$ is a function of $l_{\min}$. $l_{\min}$ that minimizes $D\left(l_{\min},\hat{l}_{\max}\right)$ is numerically chosen from the observed data. That is, $\hat{l}_{\min}=\arg\min_{l_{\min}} D\left(l_{\min},\hat{l}_{\max}\right)$. From the above equations, $\hat{l}_{\min}$ and $\hat{l}_{\max}$ are obtained. Finally, the exponent $\hat{\eta}=\hat{\eta}\left(\hat{l}_{\min},\hat{l}_{\max}\right)$ of the TP model that best fits the data in the range $\hat{l}_{\min} \le l \le \hat{l}_{\max}$, is determined using Equation (S3).

### Fitting to EP

In this section, the aim is to determine the minimum value $\hat{l}_{\min}$ of the observed data to be fitted to the EP model and the exponent $\hat{\lambda}$ of the EP model that best fits the data in the range $\hat{l}_{\min} \le l$. The EP in the range $l_{\min} \le l$ is expressed by the following equation:

$$p\left(l;\lambda,l_{\min}\right)=\lambda e^{-\lambda\left(l-l_{\min}\right)} \qquad \text{(S5)}$$

The CCDF of $p\left(l;l_{\min},\lambda\right)$ is expressed as follows:

$$P\left(l;\lambda,l_{\min}\right)=e^{-\lambda\left(l-l_{\min}\right)} \qquad \text{(S6)}$$

If the data in the range of $l_{\min} \le l$ are $\left\{l_1,l_2,\cdots,l_m\right\}$, then the log likelihood for these data is expressed as

$$L\left(\lambda;l_{\min}\right)=\sum_{i=1}^{m}\ln p\left(l_i;\lambda,l_{\min}\right)=m\ln\lambda-\lambda\sum_{i=1}^{m}\left(l_i-l_{\min}\right) \qquad \text{(S7)}$$

The exponent $\hat{\lambda}\left(l_{\min}\right)$ that maximizes $L\left(\lambda;l_{\min}\right)$ is obtained as a solution to $\frac{\partial L\left(\lambda;l_{\min}\right)}{\partial\lambda}=0$ using the following formula:

$$\hat{\lambda}\left(l_{\min}\right)=m\left(\sum_{i=1}^{m}\left(l_i-l_{\min}\right)\right)^{-1} \qquad \text{(S8)}$$

The Kolmogorov-Smirnov statistic $D\left(l_{\min}\right)$ to measure the closeness of CCDF $S\left(l;l_{\min}\right)$ obtained from the data in the range $l_{\min} \le l$ and the theoretical CCDF $P\left(l;\hat{\lambda}\left(l_{\min}\right),l_{\min}\right)$ represented by Equation (S6).

$$D\left(l_{\min}\right)=\max_{l_{\min}\le l}\left|S\left(l;l_{\min}\right)-P\left(l;\hat{\lambda}\left(l_{\min}\right),l_{\min}\right)\right| \text{(S9)}$$

$\hat{l}_{\min}$ is calculated from $\hat{l}_{\min}=\arg\min_{l_{\min}} D\left(l_{\min}\right)$. The final value is $\hat{\lambda}=\hat{\lambda}\left(\hat{l}_{\min}\right)$.

### Comparison of TP and EP

In this section, the authors describe a method for determining the most suitable distribution model (TP or EP) for the simulation data. Akaike information criteria weights (AICw) are used for comparison [6]. First, the Akaike information criterion (AIC) for data in the range $l_{\min} \le l \le l_{\max}$ is defined as follows:

$$AIC_{TP} = -2\ln\left(L\left(\hat{\eta}; l_{\min}, l_{\max}\right)\right) + 2$$
$$AIC_{EP} = -2\ln\left(L\left(\hat{\lambda}; l_{\min}\right)\right) + 2 \quad \text{(S10)}$$

The AIC difference $\Delta$ is thereafter calculated as follows:

$$AIC_{\min} = \min\left(AIC_{TP}, AIC_{EP}\right)$$
$$\Delta_{TP} = AIC_{TP} - AIC_{\min} \quad \text{(S11)}$$
$$\Delta_{EP} = AIC_{EP} - AIC_{\min}$$

Finally, AICw are calculated as follows:

$$w_{TP} = \frac{e^{-\Delta_{TP}/2}}{e^{-\Delta_{TP}/2} + e^{-\Delta_{EP}/2}}$$
$$w_{EP} = \frac{e^{-\Delta_{EP}/2}}{e^{-\Delta_{TP}/2} + e^{-\Delta_{EP}/2}} \quad \text{(S12)}$$

First, using the data in the range $\hat{l}_{\min} \le l \le \hat{l}_{\max}$ calculated during the fitting of TP, the most appropriate exponents, $\hat{\eta}$ and $\hat{\lambda}$, are determined for each model.

Next, these exponents are used to calculate and compare AICw. Subsequently, we changed $\hat{l}_{\min}$ to the one calculated during the fitting of the EP and compared. If $w_{TP} > w_{EP}$ for both data, TP is considered to fit the simulated data better. However, if $w_{TP} < w_{EP}$ for both data, EP is considered to fit the simulated data better. In case of discrepancies between the results in both datasets, the following indicators are defined and examined according to reference [2].

$$D_{adj,TP} = \frac{\ln N}{\ln n_{TP}} D_{TP}$$
$$D_{adj,EP} = \frac{\ln N}{\ln n_{EP}} D_{EP} \quad \text{(S13)}$$

where $D_{\mathrm{TP}}$ and $D_{\mathrm{EP}}$ are the Kolmogorov-Smirnov statistics calculated during the model fitting of TP and EP, respectively. $N$ is the total number of observed data points, and $n_{TP}$ and $n_{EP}$ are the number of observed data points used in each model fitting. In other words, the index considers a model that can fit more observational data. In the case of $D_{adj,TP} < D_{adj,EP}$, TP is considered to fit the simulation data better. Conversely, when $D_{adj,TP} > D_{adj,EP}$, EP is considered to fit the simulation data better.
